# Self-generation and sound intensity interactively modulate perceptual bias, but not perceptual sensitivity

**DOI:** 10.1101/2020.11.23.393785

**Authors:** Nadia Paraskevoudi, Iria SanMiguel

## Abstract

The ability to distinguish self-generated stimuli from those caused by external sources is critical for all behaving organisms. Although many studies point to a sensory attenuation of self-generated stimuli, recent evidence suggests that motor actions can result in either attenuated or enhanced perceptual processing depending on the environmental context (i.e., stimulus intensity). The present study employed 2-AFC sound detection and loudness discrimination tasks to test whether sound source (self- or externally-generated) and stimulus intensity (supra- or near-threshold) interactively modulate detection ability and loudness perception. Self-generation did not affect detection and discrimination sensitivity (i.e., detection thresholds and Just Noticeable Difference, respectively). However, in the discrimination task, we observed a significant interaction between self-generation and intensity on perceptual bias (i.e. Point of Subjective Equality). Supra-threshold self-generated sounds were perceived softer than externally-generated ones, while at near-threshold intensities self-generated sounds were perceived louder than externally-generated ones. Our findings provide empirical support to recent theories on how predictions and signal intensity modulate perceptual processing, pointing to interactive effects of intensity and self-generation that seem to be driven by a biased estimate of perceived loudness, rather by changes in detection and discrimination sensitivity.

**Highlights:**

- Self-generation and stimulus intensity interactively shape auditory perception.
- Supra-threshold self-generated sounds are perceptually attenuated.
- When near-threshold, perceived intensity is enhanced for self-generated sounds.
- Self-generation and intensity modulate perceptual bias, rather than sensitivity.
- Surprise-driven attentional mechanisms may underlie these perceptual shifts.

## 1. Introduction

The ability to make sense of the noisy information present in the world around us is crucial for our survival. Yet, what we perceive is not a veridical reproduction of the signals reaching our sensory apparatus, but it is instead an interplay between bottom-up processes and top-down predictions about the upcoming events (Friston, 2005). Forming predictions about what is about to come helps us interact with the world around us, by perceptually prioritizing behaviourally relevant sensory events. Attempts to assess how expectations influence our perception show that we are more likely to report perceiving an expected than an unexpected stimulus (Chalk et al., 2010; Jaramillo & Zador, 2011; Pinto et al., 2015; Stein & Peelen, 2015; Wyart et al., 2012). However, although the facilitatory effects of expectation on perceptual processing have been found in the wider sensory literature, they usually conflict with work from the action domain (for a recent review see Press et al., 2020).

Being able to predict the sensory consequences of our own action constitutes a specific instance of predictive processing that is highly critical in perceiving behaviourally relevant events in our environment. Several lines of research have shown that actions suppress the processing of the self-generated reafferent input (e.g., action-induced blindness, Kunde & Wühr, 2004; saccadic suppression, Ross et al., 2001; self-generation of stimuli, Straka et al., 2018). The attenuated physiological responses to self- compared to externally-generated inputs appear to be widespread throughout the animal kingdom and modality independent, being reported in a wide range of species (Chagnaud et al., 2015; Kelley & Bass, 2010; Kim et al., 2015; Requarth & Sawtell, 2011; Roy & Cullen, 2001; Schneider et al., 2014) and in several sensory modalities, including the auditory (Baess et al., 2011; Horváth, 2013a, 2013b; Martikainen et al., 2005; Mifsud et al., 2017; SanMiguel et al., 2013; Saupe et al., 2013; Schafer & Marcus, 1973; Timm et al., 2013), visual (Hughes & Waszak, 2011; Mifsud et al., 2018; Roussel et al., 2013, 2014), and tactile (Blakemore et al., 1998; Hesse et al., 2010; Kilteni et al., 2020). An influential proposal referred to as the ‘cancellation account’ attributes sensory attenuation to an efference copy of the motor command generated before or during an action that is sent from the motor to the corresponding sensory cortices (Sperry, 1950; von Holst, 1954). This efference copy allows one to accurately predict the imminent stimulation resulting from the individual’s own action via internal forward modelling (Wolpert et al., 1995). The resulting motor-driven predictions of sensory reafference (i.e., the “corollary discharge”) are then compared to the actual sensory consequences of one’s actions, and subsequently, only the difference between the two (i.e., prediction error) is sent to higher stages of the neuronal hierarchy for further processing (Friston, 2005; Wolpert & Miall, 1996), effectively cancelling out responses to predictable input. The cancelling role of the motor-driven predictions in sensory cortices has been suggested to be of great ecological importance, as it contributes in prioritizing the newsworthy unpredictable information (Barron et al., 2020), by distinguishing stimuli that correspond to potentially biologically significant external events from stimuli that arise simply as a consequence of our own motor actions (Blakemore et al., 2000; Poulet & Hedwig, 2002), and shapes our perception of sense of agency (Gallagher, 2000).

However, in the animal kingdom corollary discharge has been found to influence sensory processing in myriad ways besides cancellation of reafference (Crapse & Sommer, 2008). Contrary to cancellation theories, recent sharpening models propose that perception is biased towards the expected input (e.g., Yon & Press, 2017; Yon et al., 2020), in line with evidence showing enhanced BOLD responses to self-generated stimuli (e.g., Reznik et al., 2014; Simões-Franklin et al., 2011) and increased discharges in some neurons during self-initiated vocalizations (Eliades & Wang, 2003). The discrepancy between cancellation and sharpening accounts is also reflected in human studies attempting to assess the behavioural correlates of the neurophysiological effects of self-generation on stimulus processing. While self-initiated action effects have been typically found to be perceived as less ticklish (e.g., Blakemore et al., 1998; Claxton, 1975; Weiskrantz et al., 1971), less forceful (Bays et al., 2005; Kilteni et al., 2020), or less loud (Sato, 2008; Weiss et al., 2011a, 2011b) than equivalent stimuli initiated by another person or by a computer, recent findings show enhanced perception for action-expected outcomes (Desantis et al., 2016; Reznik et al., 2014; Yon et al., 2020). Collectively, the discrepancy in the results reported so far points to factors other than self-generation that may interactively modulate sensory processing during motor actions.

In a closer look, the mixed findings reported so far as concerns the neurophysiological and behavioural effects of motor predictions on sensory processing may be due to critical differences in the experimental paradigm, stimulus features, and obtained measures (see Table 1 for a summary of the human studies with auditory stimuli). On the one hand, animal studies with perceptual measures have reported both attenuation (McGinley et al., 2015; Neske et al., 2019) and enhancement (Carcea et al., 2017), but assess perceptual processing during locomotion compared to quiescence (Bennett et al., 2018; McGinley et al., 2015; Neske et al., 2019) or in Go compared to NoGo trials (Carcea et al., 2017). However, sensory processing during action may differ from processing of stimuli resulting from action as assessed in contingent paradigms with humans that typically compare action-predicted vs. unpredictable stimuli (i.e., self- vs. externally-generated; e.g., Sato, 2008; Kilteni et al., 2020; Weiss et al., 2011a, 2011b) or predicted vs. mispredicted stimuli (action-congruent vs. action-incongruent; e.g., Yon et al., 2020; Yon & Press, 2017), thus rendering it difficult to disentangle whether the observed effects are driven by specific motor-driven predictions or by unspecific arousal mechanisms (McGinley et al., 2015). Additionally, studies also differ in the task and stimulus intensities that they employ. Human studies reporting suppression typically use supra-threshold stimuli in discrimination paradigms and show modulations in perceptual bias (Point of Subjective Equality; PSE) rather than sensitivity measures (Just Noticeable Difference; JND, e.g., Sato, 2008; Kilteni et al., 2020; Weiss et al., 2011a, 2011b). In contrast, evidence supporting sharpening accounts has been reported mostly in detection paradigms that obligatorily need to use near-threshold stimuli (Cao & Gross, 2015; Desantis et al., 2016; Reznik et al., 2014; Yon et al., 2020; Yon & Press, 2017). This line of work has reported changes in sensitivity in both directions (e.g., Reznik et al., 2014; Cardoso-Leite et al., 2010; Cao & Gross, 2015, but see Schwartz et al., 2018 for no effects), but also in decision processes (Desantis et al., 2016; Yon et al., 2020). Collectively, these findings raise the possibility that the conflicting findings on the nature of the effects of action on the perceptual processing of self-initiated stimuli may depend on a handful of specific factors (i.e., action/no action comparisons vs. action-predicted/action-unpredicted comparisons; stimulus intensity) that may selectively affect certain aspects of perception (i.e., detection or discrimination ability; sensitivity or bias).

**Table 1.** Human studies assessing the behavioural effects of self-generation on auditory processing.

| Self-generation effects | Study | Task | Intensity | Bias / sensitivity |
| --- | --- | --- | --- | --- |
| Attenuation | Sato, 2008;<br>Weiss et al.,<br>2011a,<br>2011b | Loudness<br>discrimination | L | Bias (PSE) |
|  | Reznik et<br>al., 2015 |  |  | Bias (% 1<sup>st</sup> sound<br>louder) |
|  | Cao &<br>Gross, 2015 | Detection of<br>attended<br>frequencies | NT | Sensitivity (d') |
| Enhancement | Reznik et<br>al., 2015 | Loudness<br>discrimination | NT | Bias (% 1<sup>st</sup> sound<br>louder) |
|  | Reznik et<br>al., 2014 | Detection | NT | Sensitivity (d',<br>thresholds) |
|  | Myers et<br>al., 2020 | Loudness<br>discrimination | L | Sensitivity (% correct) |
| No effect | Sato, 2008;<br>Weiss et al.,<br>2011a,<br>2011b | Loudness<br>discrimination | L | Sensitivity (JND) |
|  | Myers et<br>al., 2020 | Detection | NT | Sensitivity<br>(thresholds) |
|  | Cao &<br>Gross, 2015 | Detection of<br>nonattended<br>frequencies | NT | Sensitivity (d') |

*Note.* Studies have reported either attenuation, enhancement, or no effects in detection or discrimination tasks with either loud (L) or near-threshold (NT) sounds by obtaining various measures that are used as a proxy of either bias or sensitivity (Point of Subject Equality, PSE; Just Noticeable Difference, JND; d', d-prime).

Recent work has indeed provided some evidence showing that sensory attenuation may be dependent on the stimulus intensity (Burin et al., 2017; Reznik et al., 2015). Reznik and colleagues (2015) had participants judge the perceived intensity of self- and externally-generated sounds presented at a supra- or a near-threshold intensity. Unbeknownst to the participants, the two sounds were always presented at the exact same intensity, but they were asked to report which one of them was louder. Their results showed a significant interaction between intensity and sound source. While the supra-threshold self-generated sounds were perceived as less loud than the passive comparisons, the opposite effect was obtained for near-threshold intensities. That is, when the sensory consequences of participants’ movements were of low intensity, a significant sensory enhancement was observed, with the self-generated tones being judged as louder than the comparison passive tones. However, due to the experimental design of this study (i.e., no varying comparison intensities), no psychophysical measures (e.g., PSE or JND) could be obtained to further examine whether the modulatory effects of intensity on perceptual processing for self-initiated sounds are driven by changes in bias or sensitivity, respectively.

Taken together, the evidence reported so far suggests that the direction of self-generation effects may be dependent on the intensity and therefore the amount of sensory noise in the signal. Indeed, recent work has highlighted the role of sensory noise in driving perceptual processing, suggesting that enhanced sensory processing for unexpected events is dependent on the ‘newsworthiness’ of the signal, such that the less the sensory noise (i.e., high intensities), the higher the sensory precision of the signal, and thus the more informative the unexpected (i.e., externally-generated) stimulus (Press et al., 2020; Barron et al., 2020). Yet, we reason that the findings obtained from the previous self-generation studies cannot provide solid conclusions on this matter, due to the use of a small range of intensities (either supra-threshold only; Sato, 2008; Weiss et al., 2011a, 2011b, near-threshold only; Reznik et al., 2014, or only one of each; Reznik et al., 2015). More importantly, the inconsistency between the studies conducted so far raises the possibility of differential effects of self-generation on different aspects of perceptual processing. Indeed, expectations have been found to yield differential effects on perceptual bias and sensitivity measures in the literature outside the action domain (e.g., Bang & Rahnev, 2017; Wyart et al., 2012). However, no systematic attempts have been made to date to assess whether motor actions alter our sensitivity to the sensory feedback or whether they result in a biased estimate of its perceived loudness.

The aim of the present study is twofold: We sought to elucidate the modulatory effects of intensity on the perceptual processing of self-generated sounds across the auditory intensity range, while systematically assessing whether the expected effects drive changes in perceptual sensitivity and/or perceptual bias. To this end, we employed a sound detection and a loudness discrimination task and compared the detection and discrimination sensitivity, as well as the possible bias in perceived loudness for self- vs. externally-generated sounds at both supra- and near-threshold intensities.

Based on previous studies with self-initiated sounds of high and low intensities, we expected to observe i) sensory attenuation for self- compared to externally-generated sounds at supra-threshold intensities and ii) sensory enhancement for self- compared to externally-generated sounds at near-threshold intensities. This interaction would be evident by better detection performance for the self- as compared to the externally-generated sounds (lower detection thresholds as in Reznik et al., 2014). Similarly, in the discrimination task, this interaction would be reflected in i) lower point of subjective equality for self-compared to externally-generated sounds at supra-threshold intensities (cf., Reznik et al., 2015; Sato, 2008; Weiss et al., 2011a) and ii) higher point of subjective equality for self- compared to externally-generated sounds at near-threshold intensities (Reznik et al,. 2015). Finally, based on previous studies reporting that self-generation only affects perceived loudness, rather than discrimination sensitivity (e.g., Sato, 2008; Weiss et al., 2011a, 2011b), we did not expect any significant differences in the just noticeable difference values, at least for the supra-threshold conditions.

The hypotheses and planned analyses for this study were preregistered on the Open Science Framework (https://osf.io/ypajr/). The Method and Results sections follow the preregistered plan.

## 2. Methods

Methods follow the preregistered plan (https://osf.io/ypajr/). The present study consisted of two two-alternative forced-choice (2AFC) tasks: a detection and a discrimination task. In the detection task, participants were presented with one sound at varying intensities and had to indicate whether it was presented in a first or a second interval of time, while in the discrimination task two sounds were presented in two different consecutive intervals of time and participants had to indicate whether the first sound (standard) or the second sound (comparison) was louder. The order of tasks was counterbalanced across participants.

### 2.1. Participants

Thirty-one healthy, normal-hearing subjects, participated in the present study. Participants were typically undergraduate university students at the University of Barcelona. Participants with hearing thresholds above 20 dB, psychiatric or neurological illness, aged below 18 or above 50 years old and who consumed drugs or pharmaceuticals acting on the central nervous system were excluded. Data from three participants (i.e., participants 2, 19, 25) had to be excluded due to technical problems or inability to comply with the task instructions, leaving data from twenty-eight participants (6 men, 22 women, *M*_*age*_ = 23, age range: 18−33 years). The sample size was defined based on the preregistered a priori power analysis. All participants gave written informed consent for their participation after the nature of the study was explained to them and they were monetarily compensated (10 euros per hour). Additional materials included a personal data questionnaire and a data protection document. The study was accepted by the Bioethics Committee of the University of Barcelona and all provisions of the Declaration of Helsinki were followed.

### 2.2 Apparatus

The visual stimuli were presented on an ATI Radeon HD 2400 monitor. The auditory stimuli were presented via the Sennheiser KD 380 PRO noise cancelling headphones. To record participants’ button presses and behavioural responses, we used the Korg nanoPAD2. The buttons of this device do not produce any mechanical noise when pressed, and, thus, do not interfere with our auditory stimuli. The presentation of the stimuli and recording of participants’ button presses and responses were controlled using MATLAB R2007a (The Mathworks Inc., 2017), and the Psychophysics Toolbox extension (Brainard, 1997; Kleiner et al., 2007; Pelli, 1997).

### 2.3 Stimuli

In the detection task we used pure tones presented binaurally with durations of 300 ms at a frequency of 1000 Hz (created using MATLAB R2007a; The Mathworks Inc., 2017). The sampling frequency was 44100 Hz, the ramp duration (duration of the onset and offset ramps) was 25 ms and a number of 16 bits per sample (cf. Reznik et al., 2014, 2015). The tone intensity ranged from 0 dB to 28 dB in steps of 4 dB for passive and active conditions.

For the discrimination task, we created pure tones with the same characteristics as those used in the detection task, except for the intensities. The intensities for the standard and comparison tones were partly based on those used in previous studies (Reznik et al., 2015; Sato, 2008; Weiss et al., 2011a, 2011b). The standard tone was always presented at a fixed intensity, while the comparison intensities varied. Specifically, the standard tones had a fixed intensity of 74 dB for supra-threshold conditions, while for the near-threshold conditions we used a fixed intensity of 5 dB above the threshold as obtained from the audiometry for the 1000 Hz sounds (cf. Reznik et al., 2015). The comparison supra-threshold stimuli varied randomly between 71 and 77 dB in steps of 1 dB, thereby resulting in seven possible comparison intensities: 71, 72, 73, 74, 75, 76, 77 (cf. Sato, 2008; Weiss et al., 2011a, 2011b). For near-threshold conditions, the comparison intensities were presented at intensities starting from 3 dB below to 3 dB above the standard intensity in steps of 1 dB, so as to match the comparison intensities of the supra-threshold conditions.

### 2.4. Procedure

Participants were seated in a soundproof chamber and auditory stimuli were presented to both ears via headphones. Visual stimuli were presented by a computer screen located in front of the participants. Prior to each task, hearing thresholds were assessed with a standard pure-tone audiometry. Additionally, practice blocks were used so that participants could familiarize themselves with each task, which also allowed us to obtain the stimulus-onset-asynchrony (SOA) between interval-cue presentation and button press in order to introduce the same visual-to-sound delay in the first passive trials.

#### 2.4.1. Detection task

Participants performed a 2-Alternative Forced Choice auditory detection task, where they had to report whether a sound of varying intensities was presented in interval one or two (Figure 1a). The sounds were either self-generated (active trials) or passively presented by the computer (passive trials).

**Figure 1.**
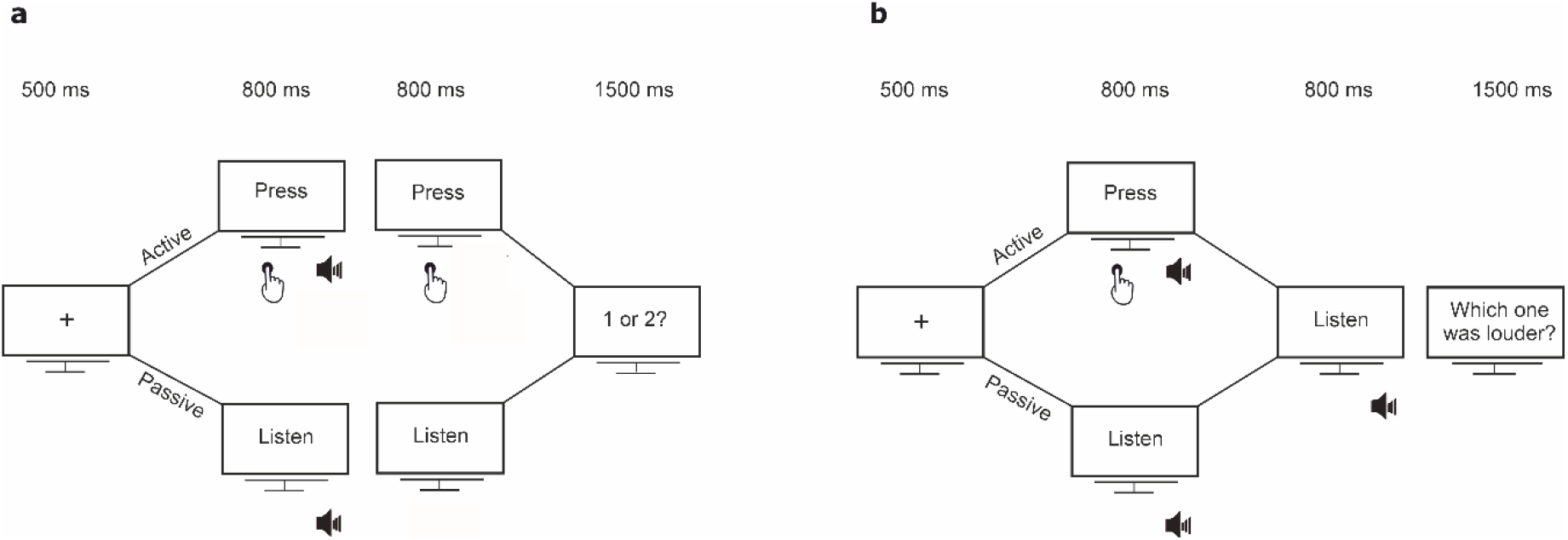
Schematic illustration of the experimental design. a) *Detection task*: Each trial started with a fixation cross, followed by two intervals. In active trials, participants were instructed to press a button in each interval (“Press” cue) and a sound was triggered either in 1^st^ or in the 2^nd^ one (in the example shown here, the sound is presented in the 1^st^ interval). In passive trials, the sound was passively presented (“Listen” cue). Participants had to respond whether they heard the sound in the 1^st^ or in the 2^nd^ interval. b) *Discrimination task*: Each trial started with a fixation cross, followed by two sounds. The first sound was either self-(active trials; “Press” cue) or externally-generated (passive trials; “Listen” cue) and was presented at an intensity of either 74 dB (suprathreshold intensity) or 5 dBs above each participant’s audiometric threshold (near-threshold intensity). The second sound was always externally-generated (“Listen” cue) and ranged ±3 dB in steps of 1 dB relative to the first one. Participants had to respond which one was louder.

Every trial started with a fixation cross with a duration of 500 ms followed by two consecutive intervals with a duration of 800 ms each. In the active trials, the sound presentation was contingent on participants’ button press. That is, participants had to press a button with their right hand once the visual cues “PRESS 1” and “PRESS 2” appeared in order to generate a sound that was triggered by the button press in either the 1^st^ or the 2^nd^ interval. For the intervals containing the sound (either 1^st^ or 2^nd^), the participants’ button press triggered the sound only if he/she pressed the button up to 300 ms prior to the interval offset. This allowed us to control that the sound had always a 300-ms duration in case a participant delayed the button press. In the passive trials, participants were passively presented with a sound in one of the two intervals indicated by the visual cues “LISTEN 1” and “LISTEN 2”. To match the timing of the sound in the active conditions, the sound was presented after an interval that was randomly selected from the participants’ distribution of press times in the active trials performed until the current trial. Thus, the timing of the stimulus presentation was equal for the two types of trials, thereby minimizing any effects of temporal predictability on the ability to detect self- and externally-generated sounds (Horvath, 2015; Hughes et al., 2013). After the offset of the second interval, the question “Did you hear the sound in the 1^st^ or 2^nd^ interval?” appeared on the screen for 1500 ms and participants had to press a button with their left hand within this time window to respond. For both trials, once a response was provided the question displayed on the screen disappeared immediately. The next trial started always after the 1500 ms response window was over.

The whole task was divided into 25 blocks consisting of 40 trials, resulting in 1000 trials in total (500 active and 500 passive trials). Active and passive conditions were presented randomly intermixed within each block (20 active and 20 passive trials). The intensities were presented using the method of constant stimuli. Intensities from 0 dB to 24 dB were presented a total of 70 times each for each condition, while we only presented the sound at 28 dB 10 times for each condition to save experimental time, given that pilot data showed ceiling performance at this intensity level. The interval containing the sound (interval 1 or 2) was random.

#### 2.4.2. Discrimination task

In the discrimination task two sounds were presented in two different consecutive intervals and participants had to indicate whether the first (standard) or the second sound (comparison) was louder (Figure 1b). Similarly to the detection task, there were two types of trials, passive and active. However, there were two additional intensity conditions, supra- and near-threshold, thereby resulting in 4 possible types of trials in total: Active and Supra-threshold (AS), Passive and Supra-threshold (PS), Active and Near-threshold (AN) and Passive and Near-threshold (PN).

Each trial started with a fixation cross with a duration of 500 ms followed by two consecutive intervals with a duration of 800 ms each. In the active trials, participants had to press a button with their right hand in the first interval, instructed by the cue “PRESS: sound 1”, in order to generate the standard tone. The comparison sound was passively presented in the second interval of time following the visual cue “LISTEN: sound 2”. The interval between visual cue and comparison sound onset was randomly selected from the participants’ distribution of press times in the first interval. For the standard self-generated sound, the participants’ button press triggered the sound only if he/she pressed the button up to 300 ms prior to the interval offset. This allowed us to control that the sound had always a 300-ms duration in case a participant delayed the button press. In the passive trials, participants were passively presented with two sounds in the 1^st^ and the 2^nd^ interval, respectively, indicated by the visual cues “LISTEN: sound 1” and “LISTEN: sound 2”. The sounds were presented after an interval that was randomly selected from the participants’ distribution of press times in the active trials. Unbeknownst to the subjects, the standard tone was always presented at the same intensity within each intensity condition: 74 dB for supra-threshold conditions and 5 dB above the threshold obtained from the audiometry for near-threshold conditions. In contrast, the comparison sound ranged from 71 dB to 77 dB in steps of 1 dB for supra-threshold conditions and ±3 dB in steps of 1 dB relative to the standard tone for near-threshold conditions. After the offset of the second comparison interval, the question “Which sound was louder: Sound 1 or Sound 2?” appeared on the screen for 1500 ms and participants had to press a button with their left hand to indicate whether the first (left button) or the second (right button) sound was louder. To control for the possibility that participants did not hear the near-threshold sounds, a third control button was used, and participants were instructed to press it only if they did not hear the two sounds. After participants’ response, the question disappeared immediately. The next trial started always after the 1500 ms response window was over.

The task was divided in 25 blocks, each one consisting of 28 trials. Each of the seven possible comparison tone intensities was presented 25 times per condition using the method of constant stimuli, as it yields a better estimation of the Point of Subjective Equality (PSE) and Just Noticeable Difference (JND) values compared to other methods (Guilford, 1954). This resulted in 175 trials per experimental condition (active/passive and supra-/near-threshold) and 700 trials in total for each participant. The conditions (i.e., sound-source: active vs. passive, and intensity: supra- vs. near-threshold) were intermixed within each block and the order of presentation was randomized for each participant.

### 2.5. Modifications from the preregistered plan

This experiment was preregistered on the Open Science Framework (https://osf.io/ypajr/). Relative to our preregistered plan, we made one modification: Instead of fitting the psychometric function with the Palamedes Toolbox (Kingdom & Prins, 2016) as reported in the preregistration of this study, we decided to use the quickpsy package in R (Linares & López-Moliner, 2016) for better visualization of the data and in order to directly introduce the values obtained from the fitting procedure to statistical analysis in R. The change in the toolbox used is not expected to have affected the results, as we kept all the parameters as predefined in the preregistration.

### 2.6. Data analysis

Data analysis follows the preregistered plan. All analysis code will be publicly released with the data upon publication (https://osf.io/ypajr/).

#### 2.6.1. Detection task

For each participant, the percentage of correct answers were calculated for each intensity and condition − active and passive −. Subsequently, for each condition, the percentage of correct responses was fitted with a normal cumulative function (Figure 2) according to the maximum likelihood procedure, using the quickpsy package in R (Linares & López-Moliner, 2016). For each participant and condition, two parameters were extracted from the model: alpha (i.e., values for thresholds in the range of the intensity levels we used) and beta (i.e., values for slope in the range of 0 to 10 in steps of .1). The lower asymptote of the psychometric function (i.e., gamma) was set to 0.5 as in previous 2-AFC detection tasks, while the upper asymptote (i.e., lambda), which corresponds to the lapse rate, was set to .001 (Kingdom & Prins, 2016). For each participant and condition, goodness-of-fit and the 95% confidence intervals for thresholds were calculated by a parametric bootstrap procedure (n = 1000; Efron & Tibshirani, 1994), using the quickpsy package in R (Linares & López-Moliner, 2016).

**Figure 2.**
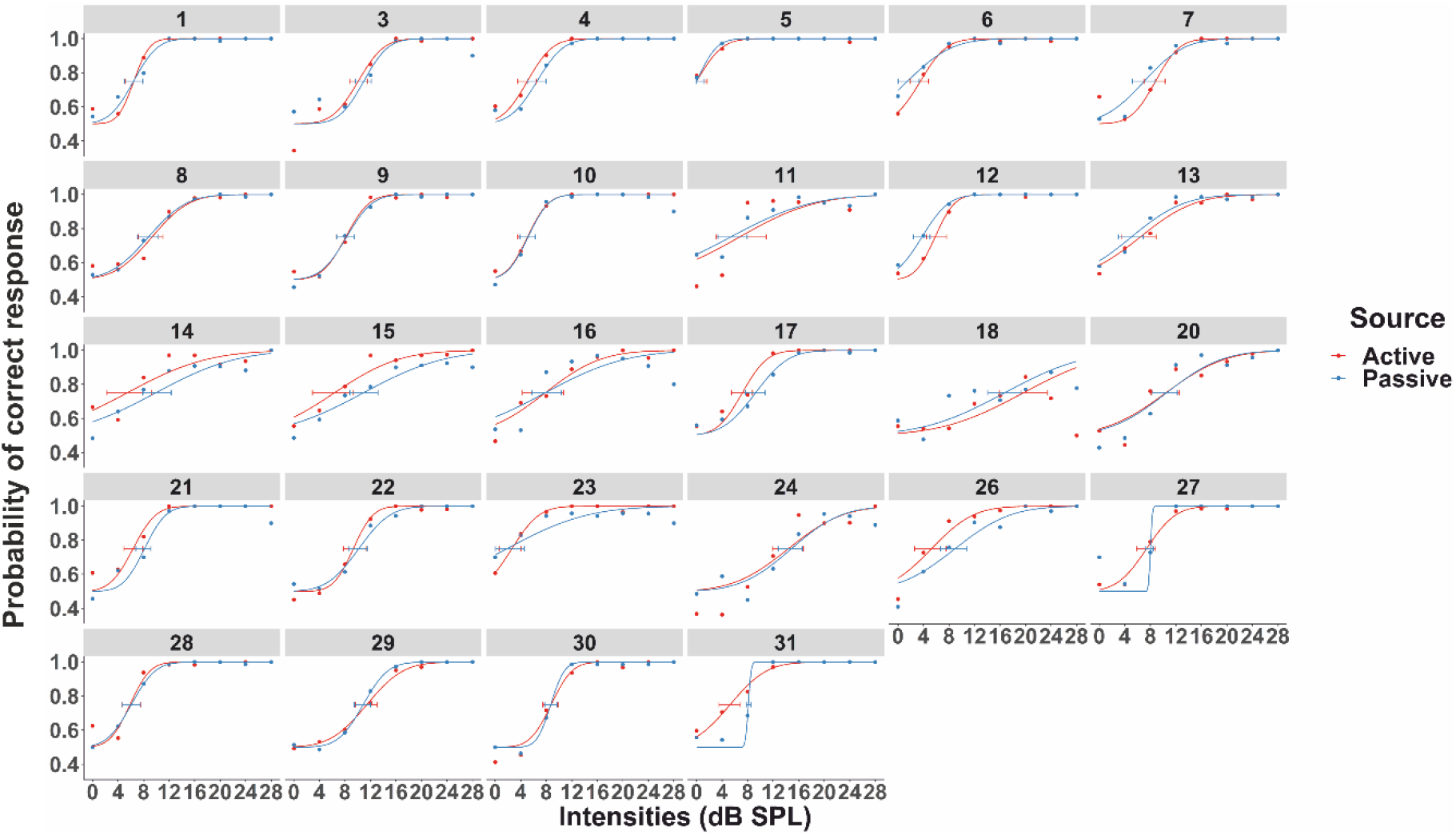
Psychometric functions for 28 participants from the detection task fitted to the percent correct responses as a function of sound intensity. Number in the legend above each plot corresponds to each participant’s number (participants with numbers 2, 19, 25 were excluded; see Methods). The small horizontal segments represent the 95% confidence intervals for thresholds (parametric bootstrap procedure with n = 1000). The threshold is defined as the intensity accurately detected at 75% of the trials (as derived from the psychometric function fitted for each participant) and is represented by the intersection of the confidence interval with the psychometric function.

The second part of the analysis consisted in calculating the d’ sensitivity index and criterion in order to directly compare our results with previous studies using this measure (Reznik et al., 2014). This analysis was performed using the Palamedes toolbox (version 1.10.3; Kingdom & Prins, 2016). Given that here we employed a 2-AFC task, we first calculated the hit and false alarm rate for one of the two intervals (interval 1 as target). As hit for interval 1 were defined the trials, where the sound was in interval 1 and the participant responded that the sound was indeed presented in this interval. As false alarm for interval 1 were defined the trials, where the participant incorrectly detected the sound in interval 1, while the stimulus was actually presented in interval 2. Subsequently, we calculated the hit rate (= number of hits divided by the number of signal trials, i.e., trials where the sound was presented in the 1^st^ interval) and the false alarm rate (= number of false alarms divided by the number of noise trials, i.e., trials where the sound was presented in the 2^nd^ interval). After z-transforming the hit and false alarm rates, we calculated the d’ (i.e., z(Hit) – z(False Alarm)) and criterion (i.e., −0.5 * z(Hit) – z(False Alarm)) for active and passive trials. Finally, we calculated the mean interval between the cue presentation and participants’ button press (henceforth SOAs) in the active trials.

#### 2.6.2. Discrimination task

For each participant, the proportion of “second sound louder” responses was calculated for each condition (active/passive, supra-/near-threshold) and for the seven comparison intensities. Data from the trials where participants did not hear the near-threshold sounds (as indicated by the third control button; see Procedure) were excluded from the analysis. In order to directly compare performance across supra- and near-threshold conditions, we defined the comparison intensities as the difference in dB from the standard stimulus: −3, −2, −1, 0, 1, 2, 3. The “second sound louder” responses for each condition were, then, fitted with a normal cumulative function (Figure 3) according to the maximum likelihood procedure, using the quickpsy package in R (Linares & López-Moliner, 2016). For each participant and condition, two parameters were extracted from the model: alpha (i.e., values in the range of the comparison intensity levels we used) and beta (i.e., values for slope in the range of 0 to 10 in steps of .1). The lower asymptote of the psychometric function (i.e., gamma) was set to 0 as in previous 2-AFC discrimination tasks, while the upper asymptote (i.e., lambda), which corresponds to the lapse rate, was set to .001 (Kingdom & Prins, 2016). Thus, for each participant and condition, two measures were obtained. First, the Point of Subjective Equality (PSE), which corresponds to the alpha values of the model, and is defined as the intensity, where the comparison stimulus was reported as louder than the standard one on 50% of the trials. This value is used to estimate the comparison tone intensity that would make the standard and comparison tones perceptually equal and is considered an index of perceptual bias (Bausenhart, Di Luca, & Ulrich, 2018). Higher PSE values would indicate that the standard first tone is perceived as louder, while lower PSE values would reflect an attenuated perceived loudness for this sound. Thus, shifts of the PSE values from the Point of Objective Equality (i.e., the point indexing the physical equality of the two sounds, which is 0 dBs here) would reflect a biased estimate of perceived loudness. Second, we extracted the just noticeable difference (JND), which corresponds to the beta values of the model (i.e., the standard deviation extrapolated from the fit) and is considered a measure of precision associated with the estimate. Higher JND values would reflect lower precision in discriminating the loudness of the two sounds (i.e., lower differential sensitivity; Gescheider, 1997). For each participant and condition, goodness-of-fit and the 95% confidence intervals for PSE were calculated by a parametric bootstrap procedure (n = 1000; Efron & Tibshirani, 1994), using the quickpsy package in R (Linares & López-Moliner, 2016). Finally, we calculated the mean interval between the cue presentation and participants’ button press (henceforth SOAs) in the active trials.

**Figure 3.**
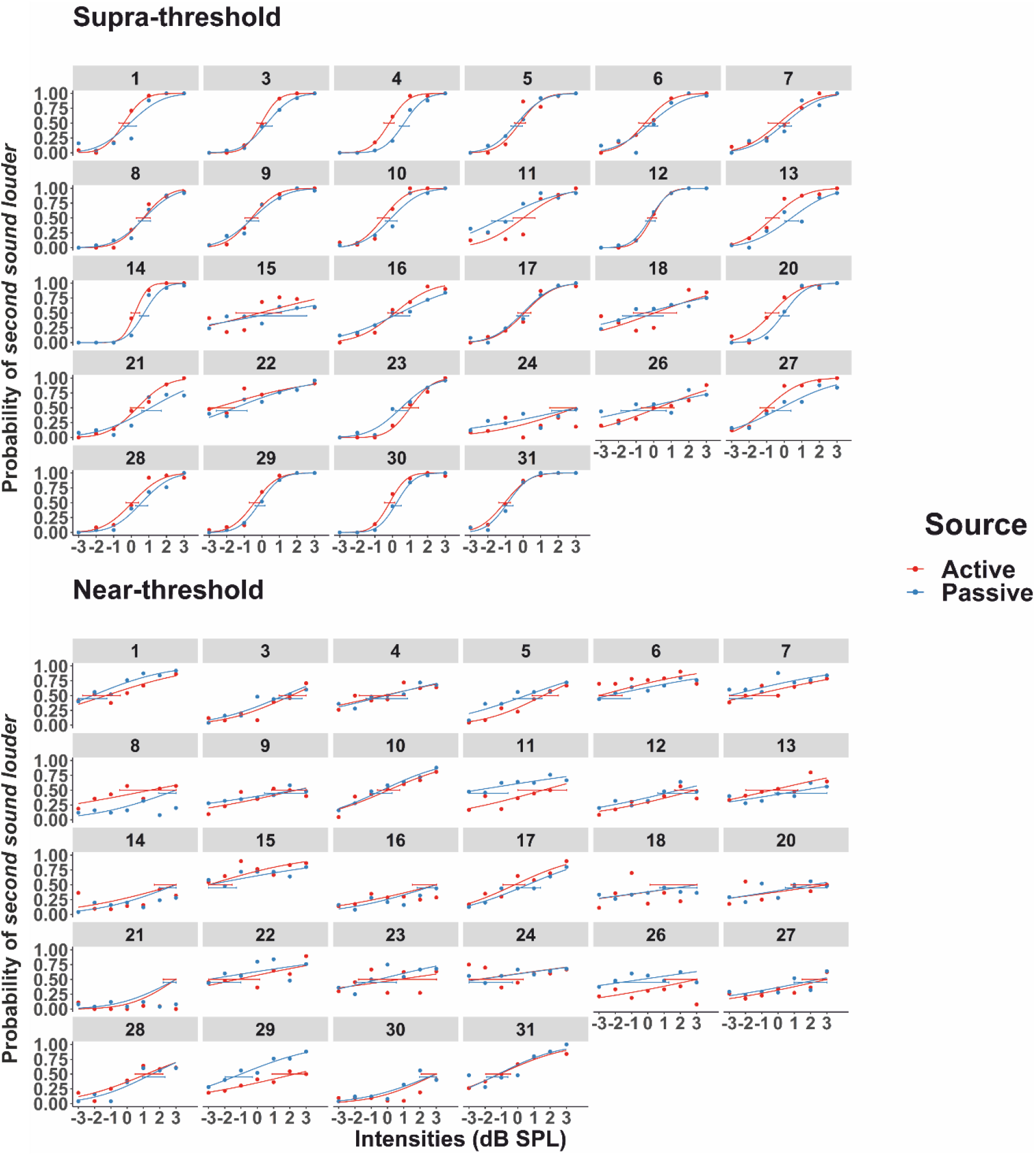
Psychometric functions for 28 participants from the discrimination task fitted to the probability of judging the comparison sound as louder as a function of its difference in dB from the first standard tone (±3 dB in steps of 1) for the supra- and the near-threshold intensities, respectively. Number in the legend above each plot corresponds to each participant’s number (participants with numbers 2, 19, 25 were excluded; see Methods). The small horizontal segments represent the 95% confidence intervals (parametric bootstrap procedure with n = 1000) for the point of subjective equality (PSE), which is defined as the intensity, where the comparison stimulus was reported as louder than the standard one on 50% of the trials.

## 3. Results

All statistical analyses were performed using R (version 3.6.0). For all the significant results in the ANOVA, we report the eta generalized squared effect size (*η_G_^2^*) and the eta partial squared (*η_p_^2^*), since the *η_G_^2^* is less biased than *η_p_^2^* (Bakeman, 2005; Olejnik, & Algina, 2003), but we also wanted to compare our findings with other studies that usually report the *η_p_^2^* effect size.

### 3.1. Modifications from the preregistered plan

This experiment was preregistered on the Open Science Framework (https://osf.io/ypajr/). Relative to our preregistered analyses, we made one modification: For the detection task, we initially planned to perform a paired-samples t-test to test for differences in the slope of the psychometric function. However, considering that the normality test was violated (Shapiro-Wilk normality test, *p* < .05), we performed a non-parametric Wilcoxon test.

### 3.2. Audiometry

From each audiometry, we obtained the thresholds for both the left and right ear. For all subjects, the thresholds were below 20 dB. Considering that in both tasks, we utilized a pure tone of 1000 Hz, in this analysis we only considered the thresholds for the 1000-Hz sounds. Specifically, for each audiometry we calculated the means across the two ears. The mean thresholds were subsequently introduced in a statistical analysis using a paired-sampled two-sided t-test to test for differences in audiometric thresholds prior to each task. The analysis did not show any significant differences (*M*_*AM_Detection*_ = 12.26, *M*_*AM_Discrimination*_ = 11.22, *SD*_*AM_Detection*_ = 3.74, *SD*_*AM_Discrimination*_ = 3.81, *p* > .05; Shapiro-Wilk normality test, *p* > .05).

### 3.3. Detection Task

The thresholds, slopes, d’, and criterion values, were analyzed using paired samples t-tests with the factor sound source − active (A) or passive (P). Trials with erroneous presses (i.e., late onset time of button press and no presses) were excluded from all analyses (*M_A_*= 28.26 %, *SD*_*A*_ = 20.37 *M*_*P*_ = 2.35 %, *SD*_*P*_ = 3.3). For the active trials, the mean interval between cue onset and button press was 0.39 s (*SD* = .07) for Interval 1 and 0.16 s (*SD* = .14) for Interval 2.

First, we performed statistical analyses for the measures obtained from the psychometric fitting procedure (Figure 4). To test for differences between the thresholds in the active and passive conditions, we used a paired samples one-tailed t-test with the hypothesis of expecting lower detection thresholds (i.e., better detection ability) in the active compared to passive trials (cf. Reznik et al., 2014; Shapiro-Wilk normality test *p* > .05). The analysis did not show any significant differences between the active and the passive conditions (*t*(27) = −1.09, *p* > .05, *M*_*A*_ = 7.46, *M*_*P*_ = 7.85, *SD*_*A*_ = 3.7, *SD*_*P*_ = 3.66), suggesting that self-generation does not have any effect on participants’ detection thresholds. Subsequently, we tested for possible differences in the slope of the psychometric function. Considering that the assumption of normality was violated (Shapiro-Wilk normality test *p* = .02), we performed a nonparametric Wilcoxon’s signed rank test for paired data on the beta values obtained from the fitting of the psychometric functions. The analysis did not show any significant difference between the active and passive slopes (*W* = 146, *p* > .05, *M*_*A*_ = 4.65, *M*_*P*_ = 5.05, *SD*_*A*_ = 2.48, *SD*_*P*_ = 3.11).

**Figure 4.**
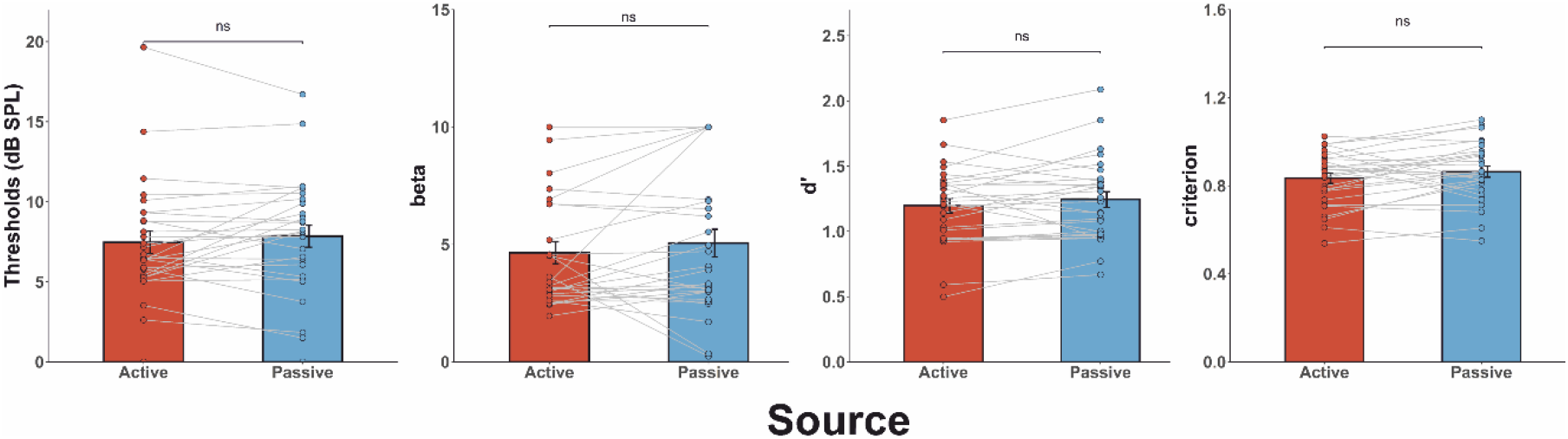
Summary of the results from the detection task. Mean (±s.e.m.) threshold, beta value for slope, d’ score, and criterion. There were no significant differences between active and passive in the threshold (one-tailed paired samples t-test, *p* > .05), slope (i.e., beta values from the psychometric fitting procedure; nonparametric Wilcoxon test due to violation of normality assumption, *p* > .05), d’ score (one-tailed paired samples t-test, *p* > .05), or criterion (two-tailed paired samples t-test, *p* > .05).

To analyze the differences in the thresholds between the two conditions, we also calculated a 95% confidence interval for the difference in thresholds based on the simulations from the bootstrapping procedure (n = 1000). For 23 out of the 28 subjects no significant differences were observed between the active and the passive trials. For one of them, the comparison between observed and simulated thresholds showed a significantly higher threshold for the active compared to the passive trials, while for the other four, a significantly lower threshold was obtained for the active trials. The goodness-of-fit routine showed that for the active trials, 26 out of the 28 psychometric curves resulted in acceptable goodness-of-fit statistics, while the fitting procedure for the passive trials showed acceptable goodness-of-fit statistics for 25 out of the 28 psychometric curves.

Subsequently, we performed a signal detection analysis for the d’ and criterion values, after confirming that the normality assumption was not violated (Figure 4; Shapiro-Wilk normality test, *p* > .05). The d’ values were analyzed using a paired samples one-tailed t-test with the hypothesis of expecting higher d’ in active compared to passive trials (cf. Reznik et al., 2014). Contrary to previous work (Reznik et al., 2014), the analysis did not show any significant differences between the active vs. passive d’ values (*M*_*A*_ = 1.2, *SD*_*A*_ = 0.3, *M*_*P*_ = 1.24, *SD*_*P*_ = .32, *p* > .05). Similarly, the criterion values were analyzed using a paired-samples two-tailed t-test (cf. Reznik et al., 2014). Similar to the findings obtained by Reznik et al. (2014), we did not observe any significant difference in the criterion values between active and passive trials (*M*_*A*_ = .83, *SD*_*A*_ = .12, *M*_*P*_ = .86, *SD*_*P*_ = .13, *p* > .05). Collectively, these findings suggest that self-generation does not affect detection sensitivity nor response bias in a 2-AFC detection task.

To further test for possible effects of self-generation and intensity level on detection performance, we also analyzed the percent of correct responses for both the active and passive trials for each one of the intensity levels. These analyses were performed so as to examine whether detection accuracy for self-generated sounds varied across the intensity levels used. We, first, conducted a repeated measures ANOVA with factors Intensity (0, 4, 8, 12, 16, 20, 24, 28) and Source (active and passive) on accuracy. The Greenhouse-Geisser correction was applied where sphericity was violated. The analysis did not show any significant main effect of source (*F*(1,27) = 1.64, *p* > .05), but we obtained a significant main effect of intensity, *F*(2.35,63.44) = 228.79, *p* < .001, *η_p_^2^* = .89 and *η_G_^2^* = .78. Specifically, irrespective of whether the sound was self- or externally-generated, participants’ accuracy was significantly lower at 0 dBs compared to the rest of the intensities, at 4 dBs compared to the intensities above 8 dBs, at 8 dBs compared to the intensities above 12 dBs, and at 12 dBs compared to intensities above 16 dBs (all *p* < .001; *M*_*0*_ = 54.65, *SD*_*0*_ = 8.85, *M*_*4*_ = 61.19, *SD*_*4*_ = 11.9, *M*_*8*_ = 77.99, *SD*_*8*_ = 13.33, *M*_*12*_ = 92.53, *SD*_*12*_ = 8.88, *M*_*16*_ = 96.47, *SD*_*16*_ = 6.03, *M*_*20*_ = 97.4, *SD*_*20*_ = 4.36, *M*_*24*_ = 97.37, *SD*_*24*_ = 4.79, *M*_*28*_ = 97.26, *SD*_*28*_ = 8.09). Comparisons between higher intensities (i.e., 16 – 28 dBs) did not show any significant differences in participants’ accuracy. The interaction between source and intensity did not reach significance (*F*(4.10,110.68) = .62, *p* > .05).

### 3.4. Discrimination Task

The PSE and JND values were analyzed using a repeated measures analysis of variance (ANOVA) with two factors: sound source − active (A) or passive (P) − and sound intensity − supra-(S) or near-threshold (N) intensity −. Trials with erroneous presses (i.e., late onset time of button press and no presses) were excluded from all analyses (*M* = 12.16%, *SD* = 10.24). For the active trials, the mean interval between cue onset and button press was 0.37 s (*SD* = .06).

The analysis for the PSE values revealed that there was not a main effect of source (*F*(1,27) = .8, *p* > .05; *M*_*A*_ = .39, *M*_*P*_ = .25, *SD*_*A*_ = 1.65, *SD*_*P*_ = 1.65) nor a main effect of intensity (*F*(1,27) = 2.62, *p* > .05; *M*_*N*_ = .65; *M*_*S*_ = -.008, *SD*_*N*_ = 2.12, *SD*_*S*_ = .86). However, there was a significant interaction between source and intensity (*F*(1,27) *=* 12.10, *p* = .002, *η_p_^2^* =.31 and *η_G_^2^ =* .15; Figure 5). The Bonferroni corrected post-hoc tests revealed a significantly higher PSE for the AN condition compared with the AS condition (*M*_*AN*_ = .92, *M*_*AS*_ = -.13, *SD*_*AN*_ = 2.04, *SD*_*AS*_ = .9, *t*(27) = −2.48, *p* = .02, *d* = .47; two-tailed post-hoc t-test), a significantly lower PSE for the AS compared to the PS condition (*M*_*AS*_ = -.13, *M*_*PS*_ = .12, *SD*_*AS*_ = .9, *SD*_*PS*_ = .83, *t*(27) = −2.41, *p* = .02, *d* = .45; one-tailed post-hoc t-test), and a significantly higher PSE for the AN compared to the PN condition (*M*_*AN*_ = .92, *M*_*PN*_ = .39, *SD*_*AN*_ = 2.04, *SD*_*PN*_ = 2.19, *t*(27) = 2.09, *p* = .02, *d* = .39; one-tailed post-hoc t-test). The post-hoc analysis did not show significant differences between the PS and the PN condition (*M*_*PS*_ = .12, *M*_*PN*_ = .39, SDPS = .83, *SD*_*PN*_ = 2.19, *t*(27) = -.64, *p* > .05; two-tailed post-hoc t-test). Thus, we replicate the findings obtained by previous discrimination studies with supra-threshold sounds (e.g., Sato, 2008; Weiss et al., 2011a, 2011b), by showing that self-generated supra-threshold sounds are attenuated compared to identical, yet externally presented stimuli. More importantly, though, we extend previous work, by showing that self-generation yields the opposite effect on perceptual bias when stimuli are presented at near-threshold intensities. That is, self-generated near-threshold sounds are perceived louder compared to the passively presented ones.

**Figure 5.**
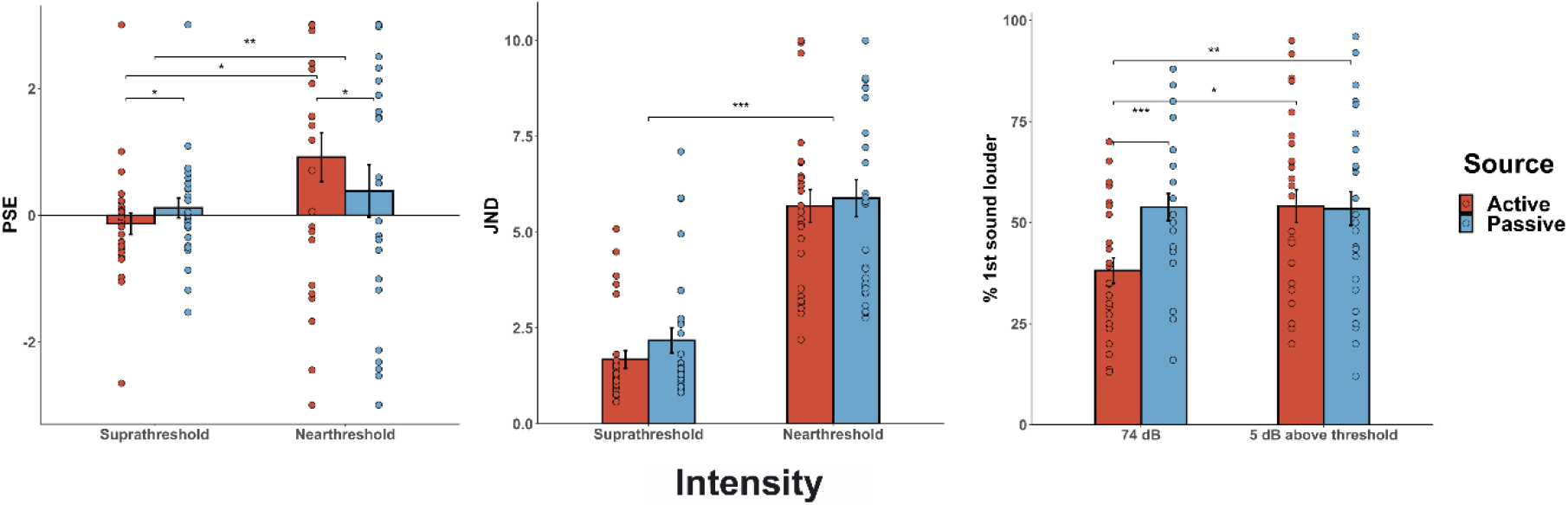
Summary of the results from the discrimination task. Mean (±s.e.m.) PSE, JND, and percent of “1^st^ sound louder responses” (cf. Reznik et al., 2015). From left to right: Significant interaction between source and intensity on PSE (*p* < .01), with the post-hoc comparisons showing lower PSE for the active supra-threshold compared to the passive supra-threshold condition (one-tailed paired samples post-hoc t-test; *p* < .05), significantly higher PSE for the active near-threshold compared to the passive near-threshold condition (one-tailed paired samples post-hoc t-test; *p* < .05), and significantly higher PSE for the active near-threshold compared to active supra-threshold (two-tailed paired samples post-hoc t-test; *p* < .05). Significant main effect of intensity on JND, with higher JND for the supra-compared to the near-threshold condition (*p* < .001). For the “1^st^ sound louder responses”, we only included trials where the standard and the comparison sounds were presented at the same intensity (i.e., 74 dB as a supra-threshold intensity and 5 dB above each participant’s threshold as a near-threshold intensity; cf. Reznik et al., 2015). There was a significant interaction between source and intensity (*p* < .01), with the post-hoc comparisons showing less “1^st^ sound louder” responses for active compared to passive trials when the sound was presented at 74 dB (one-tailed paired samples post-hoc t-test; *p* < .001), less “1^st^ sound louder” responses for active trials when presented at 74 dB compared to when presented at 5 dB above each participant’s threshold (*p* < .05), and no differences between active and passive trials when the sounds were presented at 5 dB above each participant’s threshold (one-tailed paired samples post-hoc t-test; *p* > .05).

The analysis for the JND values revealed that there was a significant main effect of intensity (*F*(1,27) = 119.45, *p* < .001, *η_p_^2^* =.82 and *η_G_^2^ =* .49), with a significantly higher JND for the supra-compared to the near-threshold conditions (*M*_*S*_ = 1.93, *M*_*N*_ = 5.79, *SD*_*S*_ = 1.5, *SD*_*N*_ = 2.39), thus pointing to lower discrimination sensitivity for near- compared to supra-threshold sounds (Figure 5). The analysis did not show a significant main effect of source (*F*(1,27) = 2.75, *p* > .05; *M*_*A*_ = 3.68, *M*_*P*_ = 4.03, *SD*_*A*_ = 2.7, *SD*_*P*_ = 2.9) nor a significant interaction between source and intensity (*F*(1,27) = .77, *p* > .05). Collectively, the results obtained by these analyses are consistent with previous work with both auditory (e.g., Sato, 2008; Weiss et al., 2011a, 2011b) and tactile self-generated stimuli (e.g., Kilteni et al., 2020) and further show that rather than being dependent on participants’ differential sensitivity in discriminating the loudness of two sounds (as indexed by the JND values), the interactive effects of intensity and self-generation are mainly driven by biases in the loudness estimates (as indexed by the PSE values).

For analyzing differences in the PSE between the four conditions, the 95% confidence intervals were calculated for each condition based on the simulations from the bootstrapping procedure (n = 1000). For 9 subjects we found significant differences between the active and passive supra-threshold conditions (for 8 subjects, lower PSE in the AS compared to the PS), while for the near-threshold intensities only 3 subjects had significantly higher PSE values in the active compared to the passive condition. Within the active condition, significant differences were obtained between the supra- and near-threshold intensities for 16 subjects (for 13 subjects, lower PSE in the AS compared to the AN), while for 18 subjects we found significant differences between the passive supra- and passive near-threshold conditions (for 12 subjects, lower PSE in PS compared to PN). The goodness-of-fit routine showed that for 26, 27, 26, and 26 psychometric curves out of the 28 total curves fitted per condition, the fitting procedure resulted in acceptable goodness of-fit statistics (for the AN, AS, PN, and PS, respectively).

Finally, we aimed to directly compare our results with the findings obtained by Reznik et al. (2015), where they employed a similar discrimination task where the standard and comparison tone were always presented at the same intensity (either supra- or near-threshold). To this end, in this analysis we only included the trials where the comparison sound was presented at the same intensity as the standard one. In particular, for the supra-threshold condition we only included the trials where the comparison sounds was presented at 74 dB (i.e., same intensity as the standard supra-threshold sounds), and accordingly for the near-threshold condition we only considered trials where the comparison tone was presented 5 dBs above each participant’s audiometric threshold (i.e., as the standard near-threshold sounds). In order to directly compare with Reznik et al.’s study, we calculated the “1^st^ sound louder” instead of “2^nd^ sound louder” responses and performed a 2×2 repeated measures ANOVA with the factors sound source (active or passive) and sound intensity (supra- or near-threshold). The results showed a significant main effect of source (*F*(1,27) = 13.54, *p* < .001, *η_p_^2^* = .33 and *η_G_^2^* = .04), with less “1^st^ sound louder” responses for active compared to passive trials (*M*_*A*_ = 46.1, *SD*_*A*_ = 20.62, *M*_*P*_ = 53.63, *SD*_*P*_ = 19.86). Contrary to the results reported in Reznik et al.’s study, the main effect of intensity did not reach significance (*F*(1,27) = 3.26, *p* > .05, *M*_*N*_ = 53.77, *SD*_*N*_ = 21.57, *M*_*S*_ = 45.98, *SD*_*S*_ = 18.76). However, consistent with Reznik et al. (2015), we obtained a significant interaction between source and intensity (*F*(1,27) = 8.94, *p* < .01, *η_p_^2^* =.25 and *η_G_^2^* = .04; Figure 5). The post-hoc t-tests showed that while there were significantly less “1^st^ sound louder” responses for AS compared to PS trials (*M*_*AS*_ = 38.12, *M*_*PS*_ = 53.82, *SD*_*AS*_ = 16.56, *SD*_*PS*_ = 17.75, *t*(27) = −5.19, *p* < .001, *d* = .98; one-tailed paired samples t-test), no differences were observed between active and passive trials when the sounds were presented at near-threshold intensities (*M*_*AN*_ = 54.09, *M*_*PN*_ = 53.45, *SD*_*AN*_ = 21.43, *SD*_*PN*_ = 22.10, *t*(27) = .17, *p* = .57; one-tailed paired samples t-test). Interestingly, consistent with the results obtained for the PSE, we also observed significantly more “1^st^ sound louder responses” for the AN compared to the AS condition (*M*_*AN*_ = 54.09, *SD*_*AN*_ = 21.43, *M*_*AS*_ = 38.12, *SD*_*AS*_ = 16.56, *t*(27) = −3.03, *p* = .01, *d* = .01; two-tailed paired samples t-test), while no differences were obtained between the PS and PN conditions (*M*_*PS*_ = 53.82, *SD*_*PS*_ = 17.75, *M*_*PN*_ = 53.45, *SD*_*PN*_ = 22.10, *t*(27) = .08, *p* = .84; two-tailed paired samples t-test). Collectively, the comparison analysis we performed replicates the significant interaction reported by Reznik et al. (2015) with an even larger effect size (*η_p_^2^* =.25 here compared to *η_p_^2^* =.21 in Reznik et al.), but the follow-up analyses demonstrate that when the standard and comparison tones are presented at the same intensity, the differences between self- and externally-generated sounds are limited to supra-threshold intensities.

### 3.5. Correlations

In an attempt to further test for possible links between detection and discrimination performance for both self- and externally-generated sounds, we conducted further correlation analyses with the values obtained from each task. Specifically, for both the self- and externally-generated sounds, we performed separate Pearson correlation analyses to assess the relationship between the detection thresholds and the PSE and JND values. For both self- and externally-generated sounds, no significant correlations were found between the detection thresholds and the PSE values (all *p* > .05). However, for self-generated sounds, we obtained significant positive correlations between the detection thresholds and the JND values for both supra- and near-threshold conditions in the discrimination task (i.e., *r* = 0.4, *p* = .04, *CI* = [.03 .67] and *r* = .48, *p* = .01, *CI* = [0.13 0.72], respectively), thus pointing to a relation between detection and discrimination sensitivity. For externally-generated sounds, we obtained significant positive correlations between the detection thresholds and the JND values, but only for the sounds presented at supra-threshold intensities (*r* = .49, *p* = .01, *CI* = [.15 .73]). Collectively, these analyses suggest that for self-generated sounds, increased detection thresholds correlate with lower discrimination precision (i.e., higher JND) for the same sounds presented both at supra- and near-threshold intensities, while increased detection thresholds for externally-generated sounds are only related with the discrimination sensitivity of the same sounds presented at supra-threshold intensities.

Additionally, we performed similar correlation analyses to test for possible links between the slope of the psychometric functions in the detection task and the PSE and JND values obtained from the discrimination task. As in the previous analysis, we did not observe any significant correlations between the slopes and the PSE values, neither for the self-nor for the externally-generated sounds. However, significant correlations were obtained again between the slopes and the JND values. For the self-generated sounds, we found significant positive correlations of the slope from the detection task with both the supra- and near-threshold conditions (*r* = .38, *p* = .05, *CI* = [.002 .66] and *r* = .4, *p* = .04, *CI* = [.1 .71]). Similarly, for the externally-generated sounds, the slopes in the detection task correlated significantly with the JND values of both the supra- and near-threshold intensities (*r* = .46, *p* = .01, *CI* = [.11 .71] for both of them).

## 4. Discussion

To-date, many different models have attempted to elucidate the effects of motor acts on perceptual processing. Yet, empirical evidence as to the exact direction and nature of these effects remain mixed. We hypothesized that the mixed findings may be related to the modulatory effects of stimulus intensity and to differences regarding the exact aspect of perceptual processing that is being tested (detection or discrimination ability; sensitivity or bias measures). Here, we present a preregistered study with a priori power estimations (https://osf.io/ypajr/), where we utilized a wide range of intensities to test for possible differences between self- and externally-generated sounds in detection and discrimination ability. Contrary to previous work (e.g., Reznik et al., 2014), we did not observe enhancements in the detection sensitivity for near-threshold self-generated sounds. However, in the discrimination task we found a significant interaction between self-generation and intensity on perceptual bias (i.e., PSE) that replicates and extends previous work (Sato, 2008; Reznik et al., 2015; Weiss et al., 2011a, 2011b) by showing that perceived intensity is reduced for self-generated sounds when they are presented at supra-threshold intensities, but enhanced when presented at near-threshold intensities.

Extant models disagree about how motor predictions affect the perceptual processing for expected action consequences. On one hand, consistent with ideomotor theories proposing that we internally activate the sensory outcome of our own action (Hommel et al., 2001), dominant cancellation models in the action literature have suggested that behavioural and neurophysiological responses to expected action effects are suppressed (i.e., lower PSE value and attenuated neural response; e.g., Blakemore et al., 2000; Kilteni et al., 2020; von Holst, 1954). Such attenuation is also predicted by preactivation accounts postulating that expectations preactivate representations of the predicted effects, increasing their baseline activity, thereby rendering the actual input less discriminable from baseline and reducing detection sensitivity (e.g., Roussel et al., 2013; Waszak et al., 2012). On the other hand, according to sharpening models, the motor-driven suppression proposed by cancellation theories is limited to units tuned away from the expected input, resulting in a sharpened population response and higher signal-to-noise ratio (SNR) that ultimately boosts detection sensitivity for what we expect (Yon et al., 2020; Yon & Press, 2017). However, none of these models can account for our findings: The cancellation account would predict lower perceived intensity irrespective of signal strength, while according to the preactivation and sharpening models we should have found significant differences in detection sensitivity (lower or higher for self-generated sounds, respectively). Critically, these models cannot explain the enhanced perceived intensity for expected sounds when presented at near-threshold intensities. Although this enhancement may be partly driven by multisensory integration processes that are known to boost processing when the unimodal signal is of low strength like the near-threshold self-generated sounds (e.g., inverse effectiveness; Stein & Meredith, 1993), two recent models have raised the possibility that the signal strength interacts with motor predictions in determining whether the processing of the expected events (i.e., self-generated sounds) will be enhanced or cancelled out (Press et al., 2020; Reznik & Mukamel, 2019).

Reznik and Mukamel (2019) recently proposed that the inhibitory influence exerted by the motor cortex on auditory areas during motor acts (Schneider et al., 2018) may either dampen or enhance perceptual processing of self-generated sounds depending on the environmental context. According to their model, the motor-driven suppression of the auditory cortex (e.g., Buran et al., 2014; Carcea et al., 2017) leads to reduced activity at the population level, but also to more selective responses and thus higher SNR. Crucially, while net activity should be always reduced during motor engagement irrespective of intensity, the resulting SNR is proposed to be higher in faint compared to salient contexts. Faint stimulation is known to elicit responses only on “best-frequency” neurons, while louder stimuli also stimulate the neurons tuned to nearby frequencies (Reznik & Mukamel, 2019). Thus, Reznik and Mukamel propose that in faint contexts, the global inhibition during motor engagement may result in “best-frequency” responses only, with almost complete silence of the activity in nearby frequencies thanks to the inhibition of the spontaneous activity, relatively enhancing the sound-evoked activity compared to the background noise (Buran et al., 2014; Carcea et al., 2017).

This proposal has two important implications as concerns the consequences of motor engagement in perceptual processing: First, salient environments would be characterized by reductions in the loudness perception that are proposed to be driven by reduced population activity. Yet, no predictions are made as to whether perceived intensity for near-threshold sounds would be also attenuated or not, thus leaving unexplained our finding that perceived intensity is enhanced for self-generated near-threshold sounds. Second, the increased SNR would boost the detectability of near-threshold sounds only, since in salient contexts sensitivity is already at ceiling. The authors found support for this claim in the study by Reznik et al., (2014), where self-generation significantly enhanced sound detectability. However, this finding was not replicated in the present study.

A caveat to the model proposed by Reznik & Mukamel is that it is largely based on animal studies that compared auditory responses in active vs. passive states (i.e., locomotion vs. quiescence or Go/No-Go tasks; e.g., Buran et al., 2014; Carcea et al., 2017), rather than comparing self-vs. externally-generated sounds. It is very likely that active states and contingent action-stimulus relationships do not have the same underlying mechanisms, and that they in turn do not modulate perception in the same way. The modulations found in active states may be mostly driven by unspecific neuromodulatory processes (i.e., arousal; McGinley et al., 2015), while in the presence of a contingent action-stimulus relationship specific prediction mechanisms may dominate (i.e., corollary discharge). This critical difference may explain why we did not find any significant effects in the detection task that lacked a contingent press-sound relationship (only 50% of the presses generate a sound). However, previous detection paradigms have also reported no such enhancement (Myers et al., 2020; McGinley et al., 2015; Neske et al., 2019), thus raising the possibility that the low power of the only human study reporting lower detection thresholds for self-generated sounds (n = 10; Reznik et al., 2014) may have reduced the likelihood of their statistically significant result reflecting a true effect (Button et al., 2013). Collectively, although Reznik and Mukamel were the first to attempt to explain how sound intensity may modulate neural and behavioural responses during motor engagement, their model cannot fully explain our findings, and in particular it also cannot explain why the interactive effects we observed here are limited to perceptual bias, rather than perceptual sensitivity.

We believe that our findings can be best explained by the opposing process theory which highlights the role of signal strength in enhancing or suppressing the processing of predictable stimuli (Press et al., 2020). According to this theory, perception is in principle biased towards expected stimuli, such as self-generated and thus more predictable stimuli. However, if the presentation of an unpredicted stimulus generates a high level of surprise after the initial stages of sensory processing, then the perceptual processing of this surprising stimulus is boosted. In terms of self-generation effects, this would imply enhanced processing of externally-generated, and thus unpredictable (surprising) stimuli. Critically, however, the level of surprise is closely related to signal strength, as surprise reflects both the distance between the prior and posterior distributions, as well as their precision (Kullback-Leibler divergence, KLD; Kullback, 1959; Itti & Baldi, 2009), and weaker signals are inherently less precise. For example, the sound of a horn in the middle of the night would elicit surprise but only if it is loud, and thus clearly audible. In sum, according to this view, supra-threshold externally-generated stimuli are inherently more surprising than the self-generated ones, shifting perception toward the unexpected (i.e., enhanced perceived loudness for the externally-generated sounds at supra-threshold intensities). Conversely, when sounds are presented at a near-threshold intensity, the increased uncertainty and higher level of noise in the signal renders externally-generated sounds unsurprising and perception is shifted towards the expected (i.e., enhanced perceived intensity for the self- compared to the externally-generated sounds at near-threshold intensities). Thus, the surprise-driven mechanism operates only for highly precise and therefore task-relevant unexpected signals, triggering a process that boosts their perception by driving attention away from the consequences of self-made acts as proposed by the active inference framework (Brown et al., 2013). Therefore, the shifts in perceived intensity in either direction may be related to surprise-induced attentional mechanisms that have been suggested to modulate the precision of the prediction error, rather than the prediction error itself (Barron et al., 2020; Brown et al., 2013). Nevertheless, one would expect that this mechanism would also operate in detection paradigms, contrary to the null findings obtained in the detection task. While these findings may be due to the lack of a contingent action-sound relationship as mentioned before, an alternative explanation is that the attentional nature of these effects results in affecting certain aspects of perceptual processing.

The studies conducted so far have not systematically assessed the effects of self-generation (and their interaction with stimulus intensity) on the different perceptual measures. Discrimination studies have only reported shifts in the PSE, a measure of *perceptual bias*, while JND – a measure of *perceptual sensitivity* – remains unaffected by self-generation (Desantis et al., 2016; Kilteni et al., 2020; Sato, 2008; Weiss et al., 2011a, 2011b). Conversely, studies employing detection tasks, have typically measured perceptual thresholds or the d’ score (*perceptual sensitivity* measures), and criterion (Cardoso-Leite et al., 2010; Reznik et al., 2014), which reflects the *response bias*. Here, we provide a more complete picture of how motor actions may affect perception by having two tasks that allowed us to obtain all these measures within subjects and show that the effects of self-generation and their interaction with stimulus intensity are driven by shifts in *perceptual bias*. This is further supported by the correlation analyses across the two tasks that yielded significant correlations only between detection thresholds and JND, both being measures of *sensitivity*, but not with *perceptual bias* measures, such as PSE. Collectively, our study points to the effects being limited to *perceptual bias*, rather than *sensitivity* measures.

In sum, the present study showed that the intensity of the sensory feedback biases perception for self-initiated stimuli in a differential manner, with attenuated perceived loudness at supra-threshold intensities, but perceptual enhancement for near-threshold ones. These findings provide empirical support to the opposing process theory (Press et al., 2020) by showing that the behavioural difference between self- and externally-generated sounds interacts with the noise of the sensory outcome in driving perceptual processing. The strength of this study is that it extends previous work by demonstrating that self-generation and its interaction with intensity only affects *perceptual bias*, rather than *perceptual sensitivity* (Myers et al., 2020; Sato, 2008; Weiss et al., 2011a, 2011b) or *response bias* (Reznik et al., 2014). Although the oppossing process theory does not clarify whether expectation effects are driven by perceptual or later decisional processes (Press et al., 2020), we argue that the proposed bias in perception as a function of signal strength implies a competition between two percepts, which was only the case in the discrimination task and may point to attentional processes that are known to reverse the effects of prediction on behavioural and neural processing (Kok et al., 2012). We believe that further behavioural and neurophysiological work is required to replicate these findings, assess the neurophysiological correlates of these effects, as well as the influence of other factors such as arousal, that are also known to affect behavioural performance (Kuchibhotla et al., 2017; McGinley et al., 2015), and ultimately provide a comprehensive account of how motor predictions and signal strength shape the perception of our environment.

## CRediT authorship contribution statement

NP and ISM designed the study; NP programmed the task and collected and analyzed the data; NP and ISM wrote the manuscript; ISM supervised the project.

## Acknowledgements

This work was supported by grants from the Spanish MINECO (PSI2014­52573­P and RYC2013­12577). The authors would like to thank Dr. Joan López-Moliner for his helpful advice on the analysis of psychophysical data and the psychometric fitting procedures.

## Notes

https://osf.io/ypajr/

### Competing Interest Statement

The authors have declared no competing interest.

